# Genome compartmentalization in a host-specific fungal insect pathogen reveals a putative mating type locus on an accessory chromosome

**DOI:** 10.1101/2025.08.27.672719

**Authors:** Dinah M. Parker, Andi M. Wilson, Carolina Nogueira, Knud N. Nielsen, Lars H. Hansen, Tue K. Nielsen, Michael Habig, Henrik H. De Fine Licht

**Author notes:** Author for correspondence, Address: Section for Organismal Biology, Department of Plant and Environmental Sciences, University of Copenhagen, Thorvaldsensvej 40, 1871 Frederiksberg, Denmark.

## Abstract

The genomic diversity of many fungal pathogens is driven by rapidly evolving regions harbouring significant amounts of transposable elements. Such genomic regions often contain genes important for pathogenicity and may be sequestered from the core genome on accessory chromosomes, which are variably present in isolates of a species. In this study, we present a pangenomic analysis of a fungal insect pathogen, *Metarhizium acridum*, using seven chromosome-scale genomes from four different continents. We show that approximately 25% of the *M. acridum* pangenome is comprised of genes only present in one isolate (singletons) or more than one but not all isolates (accessory). These genes are enriched for functions in secondary metabolite production, nutrient transport, and chromosome organization. We also find evidence of functional compartmentalization of the genome, as the core genome of *M. acridum* is enriched in carbohydrate-active enzymes, while the accessory components are enriched in effectors that are located in gene-sparse regions of the genome. Furthermore, we identified the first naturally recovered accessory chromosome in *M. acridum*, which does not harbor effector or secondary metabolite proteins related to host-insect interactions but is enriched in functions related to sexual reproduction. Within the genus *Metarhizium, M. acridum* is one of few species considered to regularly reproduce sexually. We show that the accessory chromosome contains genes from both the *MAT1-1* and *MAT1-2* idiomorphs, in addition to the *MAT* locus found on the core chromosomes. Our findings suggest that the presence of this accessory chromosome may facilitate sexual reproduction, possibly even primary homothallism, in the otherwise heterothallic *M. acridum*. Overall, our study presents a putative novel mechanism whereby this fungal pathogen may acquire gene recombination and ensure maintenance of the accessory chromosome.

## INTRODUCTION

Interactions between fungal pathogens and their hosts are shaped by a constant evolutionary struggle between immune defenses and pathogenic strategies. As hosts evolve increasingly sophisticated immune responses, fungal pathogens respond by secreting effector proteins and enzymes that promote colonization and disrupt host immunity. Such co-evolutionary interactions between adapted fungal pathogens and their hosts are considered to drive rapid evolution [1]. To facilitate this, many filamentous fungal pathogens have evolved compartmentalized genomes where pathogenicity genes are localized in distinct genomic regions [2–4]. These genomic compartments are thought to promote rapid diversification of pathogenicity-related genes in response to host selection and consequently often contain extensive genetic variation between isolates [5, 6]. Much of the intra-specific diversity is evident as gene presence-absence variation among individual isolates [7–9], and accessory regions only present in some isolates often contain genes that contribute key functional traits for virulence and environmental adaptation [10]. Collectively, these accessory genes can represent a substantial proportion of the genome—for example, up to 31% in *Aspergillus fumigatus*, 20% in *Cryptococcus neoformans*, and as much as 59% in certain plant pathogens [10–12].

The compartmentalization of fungal pathogen genomes into conserved core and fast-evolving variable accessory regions are often driven by transposable elements (TEs), gene duplication events, or horizontal gene transfers (HGTs), and can be highly variable between taxa with different lifestyles [13, 14]. Although TEs can have negative fitness consequences for the fungal genome, they are also a source of genomic innovation that can promote rapid evolution [15, 16]. One solution and evolutionary strategy to balance these opposing drivers for maintaining and eliminating TEs, is to compartmentalize the genome into TE-rich accessory regions and gene-dense core regions with few repeats [2]. These accessory regions are not only enriched in typical TEs but can also be dominated by massive elements called Starships, spanning up to 700 kb, encoding tens to hundreds of genes, and capable of horizontal transfer across species in *Pezizomycotina* fungi [17–19]. Accessory regions may be embedded within core chromosomes but they can also be physically separated from the rest of the genome and form entire accessory chromosomes. Similar to *Starships,* accessory chromosomes may be horizontally transferred between fungal species and thus can significantly contribute to rapid genomic adaptation [20, 21]. For example, the transfer of accessory chromosome 14 from a tomato-infecting *Fusarium oxysporum* isolate (Fol4287) to a nonpathogenic isolate, converted the receiving isolate into a tomato-infecting pathogen [22].

In biology, the genetic diversity and evolutionary potential is also strongly influenced by modes of reproduction and sexual strategies. Many fungal pathogens can reproduce both asexually and sexually during their life cycles, and the difference in sexual strategy is genetically determined by genes present at the mating-type (*MAT*) locus. In heterothallic fungi, mating requires a physical interaction between two compatible individual isolates with opposite copies (idiomorphs) of the *MAT* locus [23]. In contrast, homothallic fungi are able to complete the sexual cycle by themselves and typically contain both *MAT1-1* and *MAT1-2* idiomorphs within a single genome and/or cell. The primary *MAT* genes are *MAT1-1-1* and *MAT1-2-1*, which encode transcription factors with an alpha-box domain and an HMG-box, respectively [24]. The diversity and population structure of these *MAT* loci implies that even if the sexual stage has never or rarely been observed, the sexual strategy of filamentous ascomycete fungi can often be inferred. Notably, the impacts of heterothallic vs homothallic mating differ, particularly in the context of genetic diversity and evolutionary potential [25]. While heterothallic mating increases the chance for genome-wide recombination between genetically unique individuals and thus the introduction of genetic variability, the chance of mating is low due to the reliance on a suitable partner. In contrast, homothallic mating occurs independently and can thus occur more frequently, but introduces far less genetic diversity [25]. This is important because the frequency of genomic recombination during sexual reproduction contributes significantly to the speed of evolutionary change in many fungal pathogens.

The entomopathogenic genus *Metarhizium* is widespread, encompassing both generalist and specialist species. Generalists, such as *M. brunneum* and *M. anisopliae*, have broad host ranges and can infect up to seven orders of insects [26]. On the other hand, specialists such as *M. rileyi* and *M. acridum* exhibit specific pathogenicity against a single order of insects, targeting Lepidoptera and Orthoptera, respectively. Comparative genomics analyses reveal key differences between these groups. Generalist species show expansions in gene families linked to pathogenesis, while specialists exhibit rapid evolution in existing proteins [26, 27]. Additionally, horizontally acquired genes from non-fungal sources with dynamic presence/absence patterns also correlate with host specificity [28], and subtilisin-like proteases specific to *Metarhizium* are predicted to have a particularly important role in lineage diversification [29]. However, genomic differentiation between generalists and specialists may be more complex than initially defined. For example, *M. humberi*, a broad-host range species, shows evidence of sexual reproduction in its genome, a trait that was previously thought to be restricted primarily to specialists [30]. These findings highlight that host adaptation in *Metarhizium* is highly dynamic and often lineage-dependent.

The fungus *M. acridum* (Driver & Milner) J.F. Bisch., S.A. Rehner & Humber (Hypocreales: Clavicipitaceae) is a specialist pathogen that exclusively infects orthopteran insects, and is genomically predicted to have the capacity for sexual reproduction [30–32]. Unlike related taxa–*M. brunneum, M. anisopliae,* and *M. robertsii–*which are known to be plant root symbionts in addition to their capacity for insect infection*, M. acridum* has only been isolated from deceased orthopteran insects and exhibits lower rhizosphere competence [33]. Due to its specificity, select *M. acridum* isolates have significant applied value as locust-targeting biological control agents and are used commercially in Australia, Africa, Asia, and the Americas [34–39]. *M. acridum* is widespread and naturally occurs across tropical and temperate regions [40, 41]. Geographically distinct *M. acridum* isolates exhibit variation in virulence, metabolic capacity, ploidy [42], and single nucleotide genomic diversity indicative of potential local adaptation to specific orthoptera hosts [43]. Despite putative coevolution between *M. acridum* and the over 26,000 extant species in the order Orthoptera [44, 45], little is known about the contribution of accessory regions to intra-specific variation in *M. acridum* and their relation to pathogenicity.

Here, we analyze the dynamics of accessory regions in *M. acridum.* To gain insights into the genomic structure, we generated chromosome-level reference genome assemblies from six geographically distinct isolates. We used a pangenomic framework together with a previously sequenced genome (CQMa102), to elucidate the genomic structure and evolution of accessory regions and mating-type loci in genetically diverse *M. acridum* isolates.

## MATERIAL AND METHODS

### DNA Extraction and Sequencing

Six isolates of *M. acridum* were obtained from the USDA-ARS Collection of Entomopathogenic Fungal Cultures (Ithaca, NY, USA; **Fig. 1a**). Conidia from single spore cultures were inoculated in 20 mL of Sabouraud Dextrose Broth (SDB: 2.5 g L^-1^ peptone, 10 g L^-1^ dextrose, 2.5 g L^-1^yeast extract) and incubated for two days on a shaker at 25±1 °C at 130 rpm prior to DNA extraction (see supplementary text for details). DNA quality and quantity was measured using a NanoDrop 2000c and a Qubit 2.0, respectively (Thermo Fisher Scientific). All isolates were sequenced using Oxford Nanopore Technologies (ONT) on PromethION and with Illumina (paired-end) sequenced on a NextSeq550 at the Section for Microbial Ecology and Biotechnology, University of Copenhagen (**Table S1**). Nanopore sequencing libraries were prepared using the Ligation Sequencing kit SQK-LSK004 and Illumina libraries were prepared using the Nextera XT kit for paired-end sequencing.

**Fig. 1:**
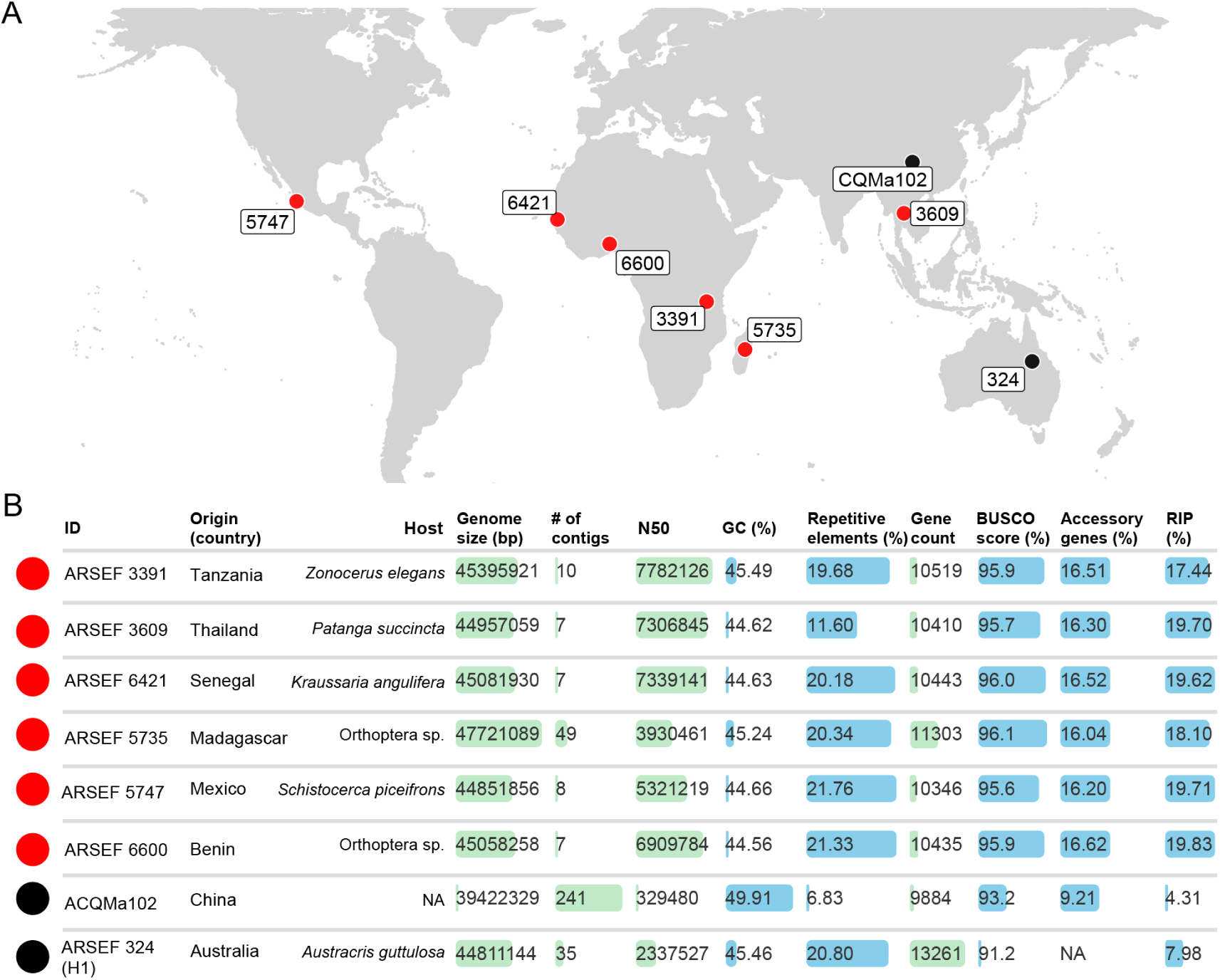
*M. acridum* isolates used for pangenome assembly. (a) Map of *M. acridum* isolates with sequenced genomes. Red points indicate genome assemblies that were generated in this study, while black points indicate genome assemblies from prior studies. (b) Isolate and genome assembly statistics for all eight *M. acridum* isolates. The bars represent the range of minimum (shortest bar) to maximum values (longest bar) for each reported statistic. Green bars indicate counts while blue bars indicate percentages. Unknown elements are indicated by NA. RIP: Percentage of genome showing evidence of Repeat Induced Point mutations, which only is active during meiosis and therefore is indicative of occasional sexual reproduction.

### Genome Assembly

First, mitochondrial ONT reads were removed if they mapped to the mitochondrial genome of *M. brunneum* isolate ARSEF 4556 (GCA_013426205; CP058939.1) [46] using minimap2 v2.17r941 [47]. Second, filtered ONT reads were assembled using both Flye v2.8 [48] and NECAT v0.01 [49]. The resulting assemblies were compared with Quast v5.0.2 [50] and the most contiguous assembly for each isolate was chosen (**Table S2**). To identify and break potential misassemblies based on long-read read coverage across the assembly, we used Tigmint v1.1.2 [51], before scaffolding with the long reads using ntLink v1.3.3 (https://github.com/bcgsc/ntLink). Detailed information is provided in the supplementary text.

### Gene prediction and annotation

Repetitive regions and transposable elements were identified using Earl Grey v5.1.1 [52] with default parameters including LTRstruct and Helitrons detection. Giant mobile elements were identified using the Starfish v1.1.0 [53] pipeline on all six genomes. Gene prediction and functional annotation of the repeat-masked polished assemblies was conducted using Funannotate v1.8.9 [54]. Protein evidence from a UniProtKB/Swiss-Prot-curated database (v2021_01) was aligned to the genomes using tBLASTn and Exonerate [55], in addition to protein evidence from *M. acridum* ARSEF 324, haplotype 1 (GCA_019434415.1) [42]. All tRNAs were predicted with tRNAscan-SE v2.0.0 [56]. Consensus gene models were found with EvidenceModeler [57]. Functional annotation were obtained using BlastP to search the UniProt/SwissProt protein database (v2021_01) [58]. Protein families (Pfam) and Gene Ontology (GO) terms were assigned with InterProScan v5.48-83.0 [59]. Signal peptides were predicted using SignalP v5.0 [60] and Phobius v1.01 [61] to identify the secretome. CAZymes were identified using HMMER v3.3 [62] and the dbCAN2 meta server [63]. Potential effectors in the secretome were identified using EffectorP v3.0^95^, putative proteases using the MEROPS database [64], and biosynthetic gene clusters using the antibiotics and Secondary Metabolites Analysis Shell v5 (antiSMASH) [65]. The completeness of the assembled genomes were evaluated using BUSCO v5.0.0 [66] with the hypocreales_odb10 data set (Creation date: 2020-08-05, 4,494 single-copy ortholog genes).

### Pan-genome and genome structure analyses

Orthologous clusters of proteins were identified using Orthofinder v2.5.4 [67, 68] with default settings using BLASTP. Orthogroup analysis included the six isolates of *M. acridum* included in this study and 16 previously published genomes of *Metarhizium* species (**Table S7)**. Orthogroups were used to generate a species tree inferred by STAG [69] and automatically rooted based on informative gene duplication events using STRIDE [70], as implemented in Orthofinder. The haplotypes of the diploid isolate, ARSEF 324 (GCA_019434415.1) were not included in the orthogroup analysis due to the nature of the un-phased genome, which confounded the results with gene duplication events [42]. Orthogroups in *M. acridum* were identified as *core* if they were present in all six isolates sequenced here and the previously published CQMa102 isolate (Fig. 1), as *accessory* orthogroups if they occurred in at least two isolates, or as *singleton* orthogroups if they only occurred in a single isolate.

Enrichment analyses of orthogroup gene ontology (GO) subsets were performed using the GOstats package in R [71], correcting for multiple comparisons via the Benjamini-Hochberg method. Orthogroups were associated with a GO term if any single isolate had a gene associated with the term. Significant differences in functional categorization of core, accessory, and singleton genome compartments were determined by Fisher’s Exact test (p < 0.05).

Local gene density was measured as 5’ and 3’ flanking distances between neighboring genes for each genome and significance was tested with a Mann-Whitney U test, with Benjamini-Hochberg-corrected p-values (p.adj < 0.05). Syntenic information was analyzed using GENESPACE v0.9.4 [72], and to perform breakpoint analysis, synteny blocks were fused if they were in the same orientation and not separated by more than five gene models. Smaller blocks entirely contained within larger blocks were removed. Breakpoint regions were defined when either the orientation or the syntenic chromosome differed, or when blocks were separated by more than five gene models. TEs overlapping these breakpoint regions were identified by comparing the locations of synteny-break regions with the TE annotation generated using the Earl Grey software pipeline, as described elsewhere, using bedtools (v2.31.1) [73]. Only DNA transposons were considered. For those at least 1,000 bp in length, the corresponding FASTA sequences were extracted and searched against the genome in which they were found using BLAST v2.16.0+. BLAST hits were filtered for at least 99% coverage and ≥90% sequence identity.

Identification of accessory chromosomes was first assessed by reciprocal cross-mapping of Illumina short reads of all six isolates using bwa-mem [74]. Second, we used pulsed field gel electrophoresis (PFGE) to assess the presence and approximate size of accessory chromosomes, analogous to the protocol described in [75]. Enrichment analyses performed on gene orthogroups located on accessory chromosomes was performed against the background of all orthogroups in *M. acridum*.

### MAT locus identification

To identify the *MAT* loci, the *SLA2* (J3458_009148), *APN2* (J3458_009151), *MAT1-2-1* (J3458_009150), and *MAT1-2-3* (J3458_009149) genes and their CDS annotations were extracted from the *M. acridum* strain ARSEF 324 H1 genome and used as queries in local BLASTn searches against the six *M. acridum* genomes using CLC Genomics V22.0. In the genomes where the *MAT1-2-1* and *MAT1-2-3* genes were not identified, the genes present in the region between the *SLA2* and *APN2* genes were confirmed as *MAT1-1-1* and *MAT1-1-3* via BLASTn and BLASTp searches against the NCBI NR database. The *MAT1-1-1* and *MAT1-1-3* genes from *M. acridum* ARSEF3391, the *MAT1-2-1* and *MAT1-2-3* genes from *M. acridum* ARSEF 5747, and the *MAT1-1-2* gene from *M. anisopliae* (NBRC 322258) as well as their protein sequences were used in BLASTn and tBLASTn searches against all six genomes to determine whether any *MAT*-like sequences were present elsewhere in the genome. Annotated *MAT* loci were compared and visualized using CLINKER as implemented in the online version of CAGECAT [76] using default parameters. For phylogenetic comparison the *MAT* loci and associated genes were identified as described above in related species, and aligned using default settings in MAFFT v7.0. The species *Pochonia chlamydosporia* and/or *Ustilaginoidea virens* were used as outgroups, depending on sequence availability. MrModelTest2 v2.4 was used to test the models for each of the alignments independently, after which Bayesian Inference (BI) analyses were conducted using MrBayes v3.2.7. The analyses were run for 1,000,000 generations, with ten parallel runs and four chains. Trees were sampled every 100 generations. Posterior probabilities were calculated from the trees that remained after 25% of the sampled trees were discarded as burn-in.

## RESULTS

### Extensive chromosomal rearrangements within the M. acridum pangenome

To provide a comprehensive view of global genomic variation in *M. acridum*, we sequenced long-read ONT libraries to a depth of 80-120X and a median read length of ∼10kb from six isolates from four continents **(Table S1, Fig 1)**. For five of the six isolates, we were able to generate chromosomal level assemblies using Flye with a stable recovery of seven chromosomes (**Fig. 1b, S1∫)**. For isolate ARSEF 5375, assemblies with both Flye and NECAT resulted in fragmented assemblies, but the latter resulted in a more contiguous assembly (49 contigs; N50=3,930,461bp; **Table S2**). Assembly sizes ranged from 44.9 Mb (ARSEF 5747) to 48.1 Mb (ARSEF 5735), and BUSCO scores were all higher than 95.6% (**Fig. S2**).

We compared whole-genome synteny across all 6 assemblies of *M. acridum* (**Fig. 2a**), which revealed that while large-scale gene order is conserved among the genomes, there is evidence for large chromosomal translocations and inversions. The positions of syntenic breakpoints in each genome were validated by Tigmint analysis, which found no evidence of breaks at these sites based on long-read sequence coverage. Two isolates displayed perfect syntenic resolution, ARSEF 6421 and ARSEF 3609, despite originating in Senegal and Thailand, respectively (**Fig 2a**). In contrast, scaffold size variation arises from translocations among ARSEF 6600, ARSEF 5747, ARSEF 3391, and the two syntenic isolates; notably, the longest scaffold is not syntenic in this four-way comparison **(Fig. 2a)**. Additionally, because *M. acridum* is known to exhibit signs of sexual reproduction, we determined that 17.5% to 19.9% of the genomes are affected by the repeat induced point (RIP) mutation processes (**Fig. 1b**).

**Fig. 2:**
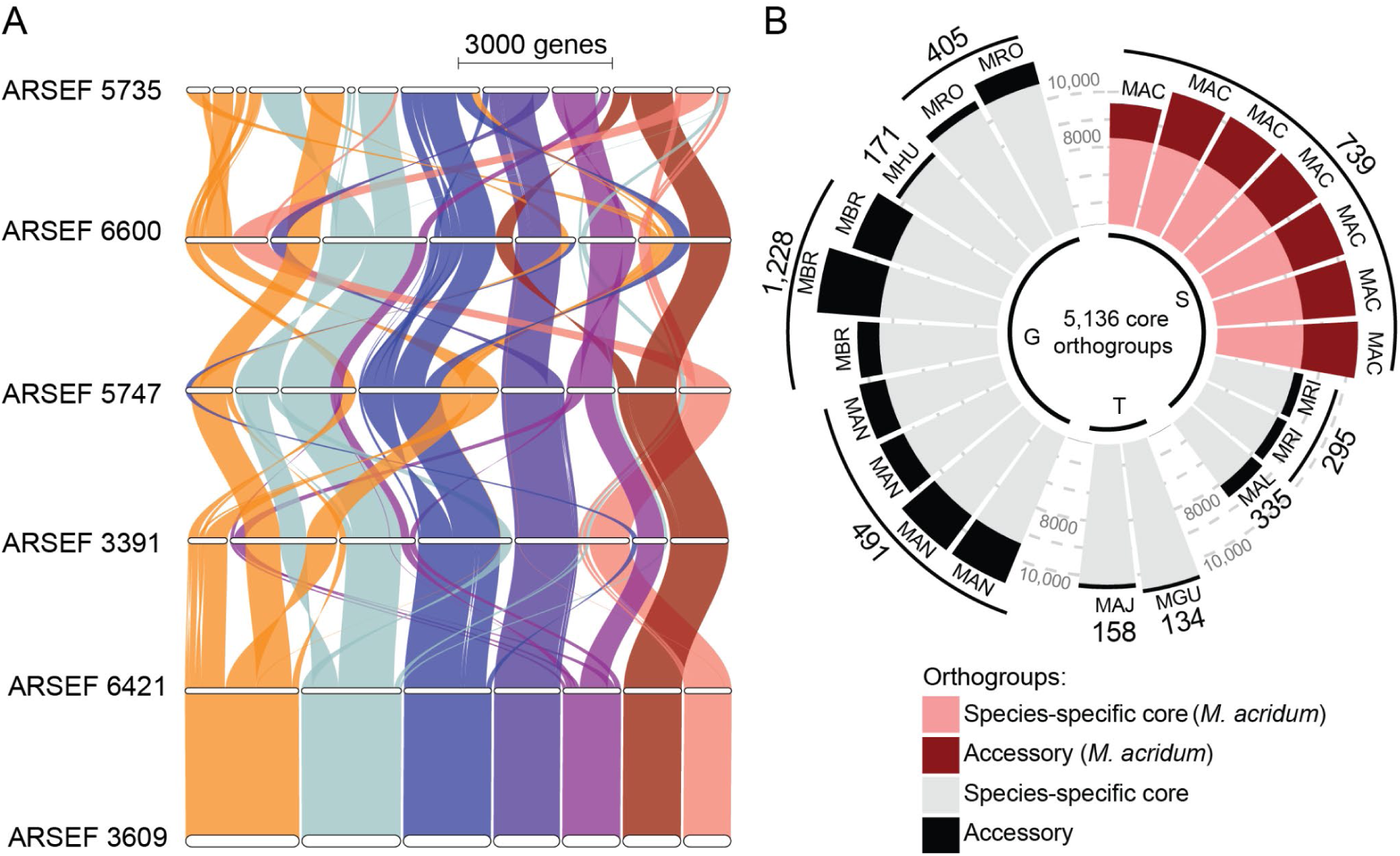
Orthogroup structure in *Metarhizium.* (A) Syntenic map of orthologous regions on core chromosome (scaffold) organization among six *M. acridum* genomes showing extensive translocation and transversions among *M. acridum* isolates. Different colours denote one of the seven core chromosomes based on ARSEF 3609. (B) Bar chart of 17,106 orthogroups found in 22 genomes of *Metarhizium*, representing 9 species. Species are indicated by abbreviations (MAC, *M. acridum*; MRI, *M. rileyi*; MAL, *M. album*; MGU, *M. guizhouense*; MAJ, *M. majus*; MAN, *M. anisopliae*; MBR, *M. brunneum*; MHU, *M. humberi*; MRO, *M. robertsii*). External values indicate the number of orthogroups (accessory and singletons) unique to each species. Insect host range is specified with G: Generalist (Up to seven insect orders), T: Transitionalist (two insect orders), and S: Specialist (a single insect order), following Hu et al. (2014).

### Transposable element (TE) divergence landscapes

To determine the repeat content of *M. acridum* isolates, we used RepeatModeler and RepeatMasker as implemented in the Earl Grey software pipeline to identify and classify repeats across genomes. We found that the six isolates of *M. acridum* contained repetitive sequences representing between 11.60% and 21.76% of the genome (mean of the genome as total interspersed repeats (**Fig. 1b**). The transposable element (TE) landscape was dominated by long terminal repeats (LTRs), which were the most abundant elements across the genomes (mean composition: 8.70% ± 0.98% SE). All six isolates showed a peak of divergence between 5-10%, suggesting an active TE expansion (**Fig. S3**). In particular, the TE landscapes of ARSEF 3391, ARSEF 5735, and ARSEF 5747 showed a greater proportion of TEs at ∼0% divergence. This suggests a more recent TE expansion in these isolates driven by DNA transposons (ARSEF 3391) or LTR’s (ARSEF 5735 and ARSEF 5747).

Recently, the horizontal acquisition and subsequent expansion of TEs were shown to result in extensive reshuffling of the genome in one strain of *M. anisopliae*, as recently acquired DNA transposons were present at many synteny breakpoints [77]. Because we observed many synteny breaks between *M. acridum* ARSEF 6421/3609 and the four other *M. acridum* isolates (**Fig. 2a**), we analyzed the breakpoints in the gene-based synteny using *M. acridum* ARSEF 6421 as reference for a putative ancestral genome organization. Within the four *M. acridum* isolates (ARSEF 3391, ARSEF 5735, ARSEF 5747, and ARSEF 6600), we found a total of 98 breakpoints compared to the ancestral state, ranging from 16 in ARSEF 5747 to 29 in ARSEF 5735 (**Table S3**). These synteny breakpoints include inversions within chromosomes or scaffolds, as well as breakpoints that connect regions originally located on two different chromosomes (**Fig. 2a**). Within these breakpoint regions, we identified 28 regions containing at least one annotated DNA TE, including 12 in *M. acridum* ARSEF 3391.

We next restricted our analysis to TE sequences at least 1,000 bp in length and assessed whether the DNA TE sequences associated with breakpoint regions had similar copies elsewhere in their respective genomes, which would indicate recent increases in copy number. Since RIP appears to be active in *M. acridum* (**Fig. 1b**), which results in rapid accumulation of mutations in repeated sequences, we used a relaxed threshold of 90% sequence identity and target sequence coverage of at least 99% of the sequence length. This revealed that in *M. acridum* ARSEF 3391 in particular, three types of DNA TEs, hAT-Ac, Helitron, and MULE-MuDR, have recently undergone copy number expansion, and these are often present at breakpoints. A total of 11 out of 27 breakpoints in ARSEF 3391 contain at least one of these recently expanded TEs (**Table S3**). While similar cases of recently expanded TEs are observed in ARSEF 5735 and ARSEF 5747, the association with breakpoints is less clear in these strains. No recently expanded DNA transposons are associated with breakpoint regions in ARSEF 6600. We therefore conclude that DNA transposons are likely involved in the genome reorganization of *M. acridum* ARSEF 3391, but it remains unclear whether this is a general feature of the species.

Annotation of giant *Starship* mobile elements, initially revealed 69 proteins with a DUF3435 tyrosine recombinase domain, which is known as the so-called ‘captain’ element of *Starships* [19]. Further visual verification of insertion positions across the *M. acridum* genomes revealed three 405,600 bp sized *Prometheus* and four ∼85,000 bp *Phoenix* family *Starship* elements (**Table S4**). The three *Prometheus* elements identified in ARSEF 6600, 6421, and 3609 share 77% of InterPro domains among their *Cargo* genes, whereas three of the *Phoenix* elements from the same three *M. acridum* isolates share 100% of InterPro domains among *Cargo* genes (**Table S5**). Analyzing InterPro domains of *Cargo* genes compared to the rest of the respective genomes, showed that *Cargo* genes are functionally enriched for carbohydrate metabolic functions but not virulence-associated genes (**Table S6**).

### Substantial gene content variation across the Metarhizium pangenome

To study variation in gene content, an orthogroup clustering analysis was performed on 16 available genomes of *Metarhizium* from 9 different species, and the 6 newly sequenced genomes of *M. acridum* generated in this study. In total, 17,106 orthogroups were identified in this genus, with 5,215(30.4%) found in all isolates, 8,793 (51.4%) occurring in at least 2 isolates, and 3,098 (18.1%) singletons (**Fig. 2b, S4**). Of the core orthogroups, 4,349 (25.4%) were defined as single copy orthologs. This confirms that even with improved genome resolution in *M. acridum* isolates, resulting in increased protein discovery compared to the previously sequenced reference (CQMa102), specialist species overall still exhibit fewer orthogroups than generalist *Metarhizium* species **(Fig. 2b)**. Additionally, the largest number of gene duplication events occurred at the node that separates the specialists (*M. acridum*, *M. rileyi*, and *M. album*) from the remaining species with larger host ranges, characterized by 103 orthogroup duplication events that are retained in at least 50% of descendent species (**Fig. S4**).

Next species-specific orthogroups were examined, defined as protein clusters that are both unique and universal to all available genomes of a given species **(Table S7)**. In *Metarhizium* species with only a single available genome, no distinction can be made between species specific orthogroups or singletons. Despite this limitation, the specialists *M. acridum* and *M. rileyi* have 255 and 343 species-specific orthogroups respectively, representing the largest collection among all sampled species. In contrast, the generalists *M. anisopliae* and *M. brunneum* only have 6 and 19 species specific orthogroups, respectively. Based on presence/absence of orthogroups, distribution of CAZymes, InterProScan, PFAM domains, or transcription factors, there is no evidence for substructure between isolates within *M. acridum* (**Fig. S4-S7**). However, three are unique combinations of orthogroups in different isolates of*M. acridum*, with the most apparent difference in ARSEF 5735, which exhibits more duplicated genes within orthogroups (**Fig. S4**). Enrichment analysis of Gene Ontology functional annotations on species-specific orthogroups in *M. acridum* showed significantly enriched categories including secondary metabolite, mycotoxin, and peptidase production, which represent enzymes known to play important roles in the infection process (**Fig. S8**). We also notably find 14 orthogroups in this subset with enriched functions in heme and tetrapyrrole binding.

### The M. acridum pangenome contains core and accessory gene compartments

To identify intraspecific gene clustering patterns in *M. acridum*, we subset orthogroups that occurred in the six *M. acridum* genomes generated in this study and the previously sequenced CQMa102. Here we identified a total of 11,046 orthogroups in *M. acridum*, with 8,331 (75.4%) core orthogroups, 2,106 (19.1%) accessory orthogroups, and 609 (5.5%) singletons (**Fig. S9a**). From the total, we determined that 7,203 (65%) were single-copy orthogroups in *M. acridum*. Singletons were found in all isolates but were more numerous in ARSEF 3391 (n = 147), ARSEF 5735 (n = 99), and CQMa102 (n = 223) (**Fig. S9b**). There was a significant difference in average protein length (aa) among the pangenome orthogroup compartments (p < 0.01), with the core compartment containing significantly longer protein-coding genes followed by the accessory and singleton compartments containing the shortest length of protein-coding genes (**Fig. S9c**).

Core orthogroups in *M. acridum* were found to be significantly enriched for “housekeeping” genes, including GO terms associated with carbon-based metabolic and catabolic processes, as well as ribosome biogenesis (**Fig. S9d**). In contrast, accessory orthogroups were found to be enriched for secondary metabolite production. Additionally, we found enrichment in functions related to chromosome rearrangement and organization, with potential associations with meiosis (**Fig. S9d**). The singleton compartments were enriched for GO terms associated with transport of glycine and amino acids (**Fig. S9d**). Overall, our results show a moderate accessory gene content in *M. acridum* (24.6%), with functions potentially related to host-pathogen interactions, metabolism, and reproduction.

To analyze how gene function was structured across the pangenome components (core, accessory, and singleton), we compared proportions of orthogroups annotated with a specific function in each component. Core orthogroups had a higher proportion of the secretome, genes with transmembrane domains, conserved protein domains, and CAZymes (**Fig. 3a-f**). In contrast, the proportion of secondary metabolite orthogroups were equal across all three pangenome compartments (**Fig. 3b**), whereas effector proteins represented the only functional category with a higher proportion in the accessory component and thus showing significant variation between individual isolates of *M. acridum* (**Fig. 3f**). A total of 289 orthogroups (2.62%) encode at least one predicted effector, and among the pangenome components, effectors made up 2.4% and 2.1% of the core and singleton orthogroups, respectively. Effectors made up 3.75% of accessory orthogroups, representing a significantly higher proportion (**Fig. 3f**).

**Fig. 3:**
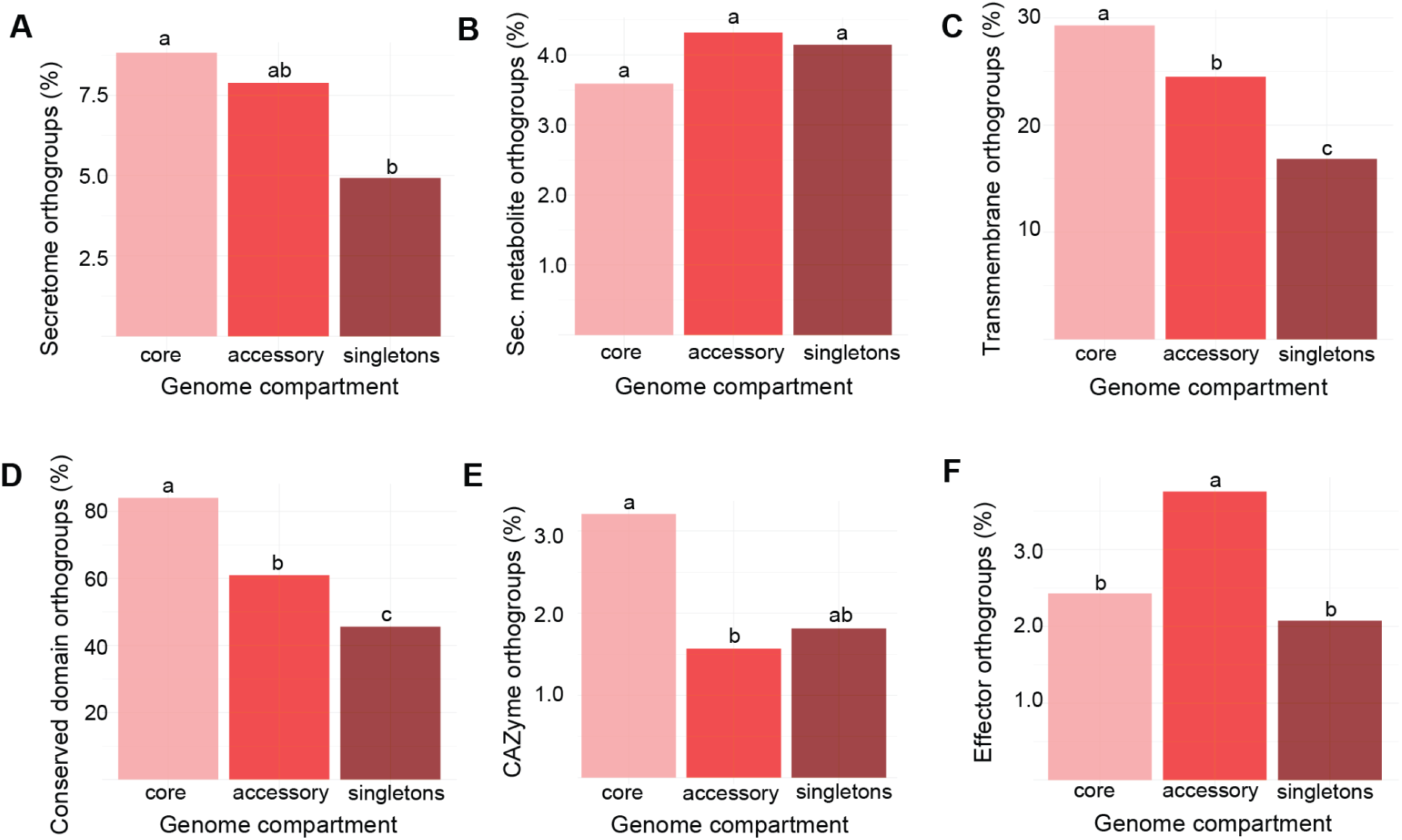
Functional categorization of *M. acridum* pangenome. Proportions of core, accessory, and singleton categories in eight functional groupings of proteins: (A) Secreted proteins, (B) secondary metabolites, (C) proteins with at least one transmembrane helix domain, (D) proteins with conserved domains, (E) CAZymes, and (F) predicted effectors. Letters indicate significance as predicted with Fischer’s Exact test. Note the different y-axis scales.

### The M. acridum genome is compartmentalized into gene-dense and gene-sparse regions

In order to evaluate if the genomes of *M. acridum* displayed genome compartmentalization, we determined the intergenic distance of each gene, and found the average distance was between 2,640 to 2,790 bp for all isolates. Density plots of the 5’ and 3’ intergenic distances showed a continuum of intergenic distances for all genomes **(Fig. 4)**. Comparison of genes with long and short intergenic distances (i.e. upper-right quadrant vs. lower-left quadrant, **Fig. 4)**, showed that the genome of *M. acridum* is compartmentalized into gene-sparse compartments with a higher density of transposable elements and gene-dense compartments with few transposable elements enriched for genes associated with fundamental molecular metabolic functions (**Fig. S10-S11)**. Considering effectors as a subset of genes, effectors occur significantly more in gene-sparse regions in all six *M. acridum* genomes (**Fig. 4a**), whereas the secretome as a whole is evenly spread across genome compartments (**Fig. 4b**). We find the same pattern for the Glycoside Hydrolase 18 (GH18) family that contains chitinase enzymes potentially used by *M. acridum* when penetrating locust cuticles [78], which are significantly more prevalent in gene sparse regions (**Fig. 4c**), whereas CAZymes as a whole are not (**Fig. 4d**). Entomopathogenic fungi also rely on Peptidase S8 as cuticle-degrading proteases [79], which are predominantly located in gene-sparse compartments in *M. acridum* (**Fig. 4e**). Although enrichment analysis showed that *M. acridum* contains a distinct set of secondary metabolite pathway genes compared to other *Metarhizium* species (**Fig. S8**), these genes are not significantly more prevalent in gene-sparse regions (**Fig. 4f**).

**Fig. 4:**
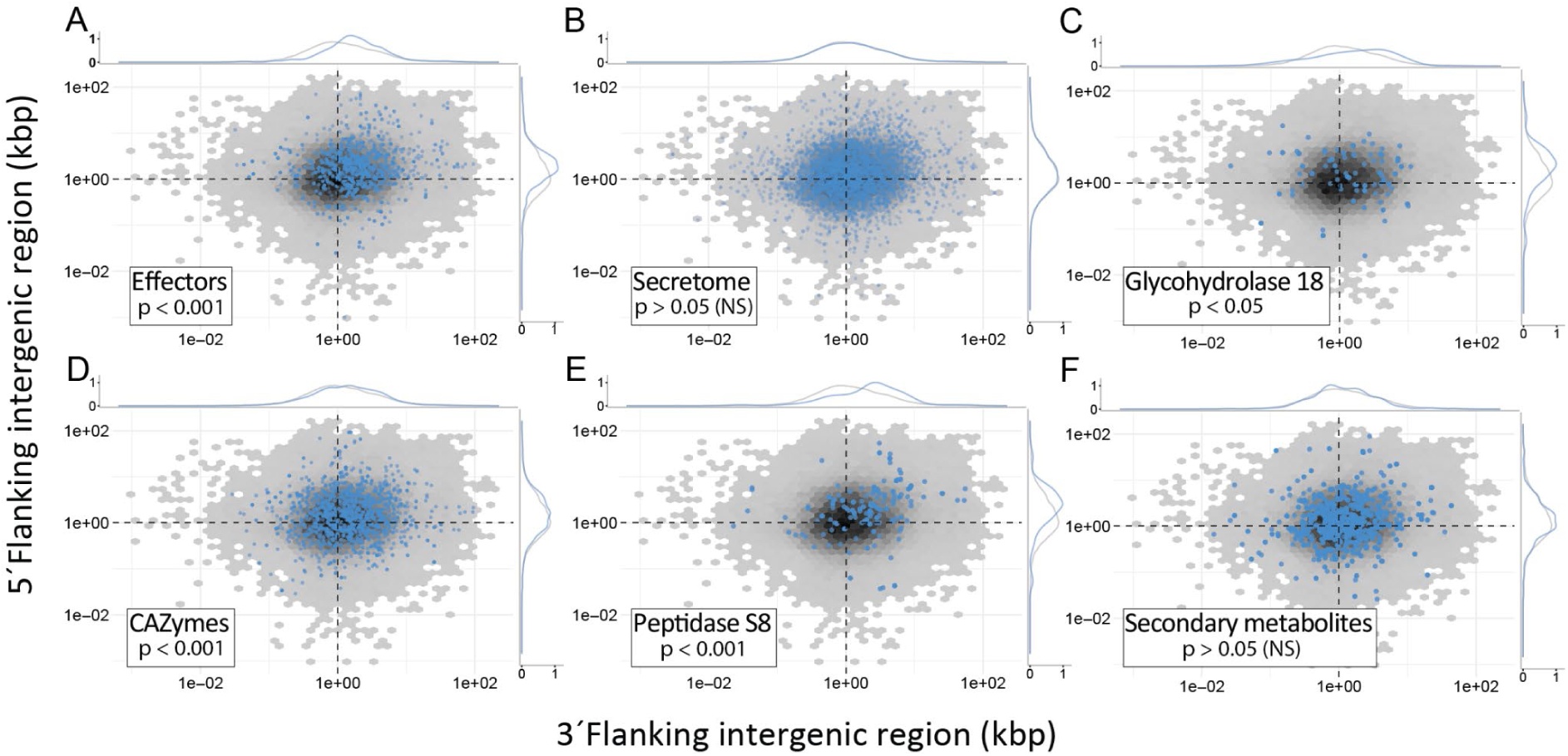
Genome compartmentalization of effectors in *M. acridum.* Gene density as a function of flanking 5’ and 3’ intergenic region sizes (x- and y-axis, respectively) of the six isolates of *M. acridum* with high-quality genome assemblies generated in this study. Hexagonal heatmaps indicate the intergenic lengths of genes with grey-to-black indicating the frequency distribution (gene count). Overlaid distribution plots in blue depict the frequency of the distributions of (A) effectors, (B) secreted proteins, (C) Glycosyl Hydrolase 18 (GH18) family enzymes, (D) CAZymes, (E) Peptidase S8, and (F) Secondary metabolites versus all other genes. P-values indicate Benjamini-Hochberg corrected values of statistical significance of intergenic lengths of gene subsets performed with the Mann-Whitney U test (NS: Not Significant).

### M. acridum contains an accessory chromosome with a putative mating type locus

We used all-vs-all cross-mapping of Illumina short reads between our six sequenced *M. acridum* isolates and observed notably lower mapping coverage of reads from other isolates for scaffolds 8 and 9 in isolate ARSEF 3391 (**Fig. 5a**). These two scaffolds each contain telomeres on one end, and to verify that scaffold 8 and 9 represent a combined accessory chromosome, we used pulsed field gel electrophoresis. This confirmed that there is an extra band only present in isolate ARSEF 3391 with a size of 1.6 Mb that nearly corresponds to the combined length of 1.5 Mb of scaffold 8 and 9 (**Fig. S12**). While the accessory chromosome has similar repeat content as core chromosomes, the gene density was lower (**Fig. 5b**). The gene-wise relative synonymous codon usage varies between the accessory and core chromosomes suggesting different evolutionary histories (**Fig. S13**). Annotation of the TE landscape per scaffold showed that TEs on the seven core chromosomes are on average made up of 52% LTR retrotransposons, whereas LTRs are not very abundant on the accessory chromosome (5% on scaffold 8 and 0.4% on scaffold 9) (**Fig. S14**). In contrast, the TEs on the accessory scaffolds contain 31% long interspersed nuclear elements (LINEs), which only constitute 0.2% of TEs on the core scaffolds. To investigate the functional role of genes on the accessory chromosome, we conducted an GO-term enrichment analysis of orthogroups present on the accessory chromosome against all *M. acridum* isolates. This revealed several enriched categories associated with functions related to sexual reproduction, such as mating type determination, cell fate commitment, and mating type specific transcription (**Fig. S15**).

**Fig. 5:**
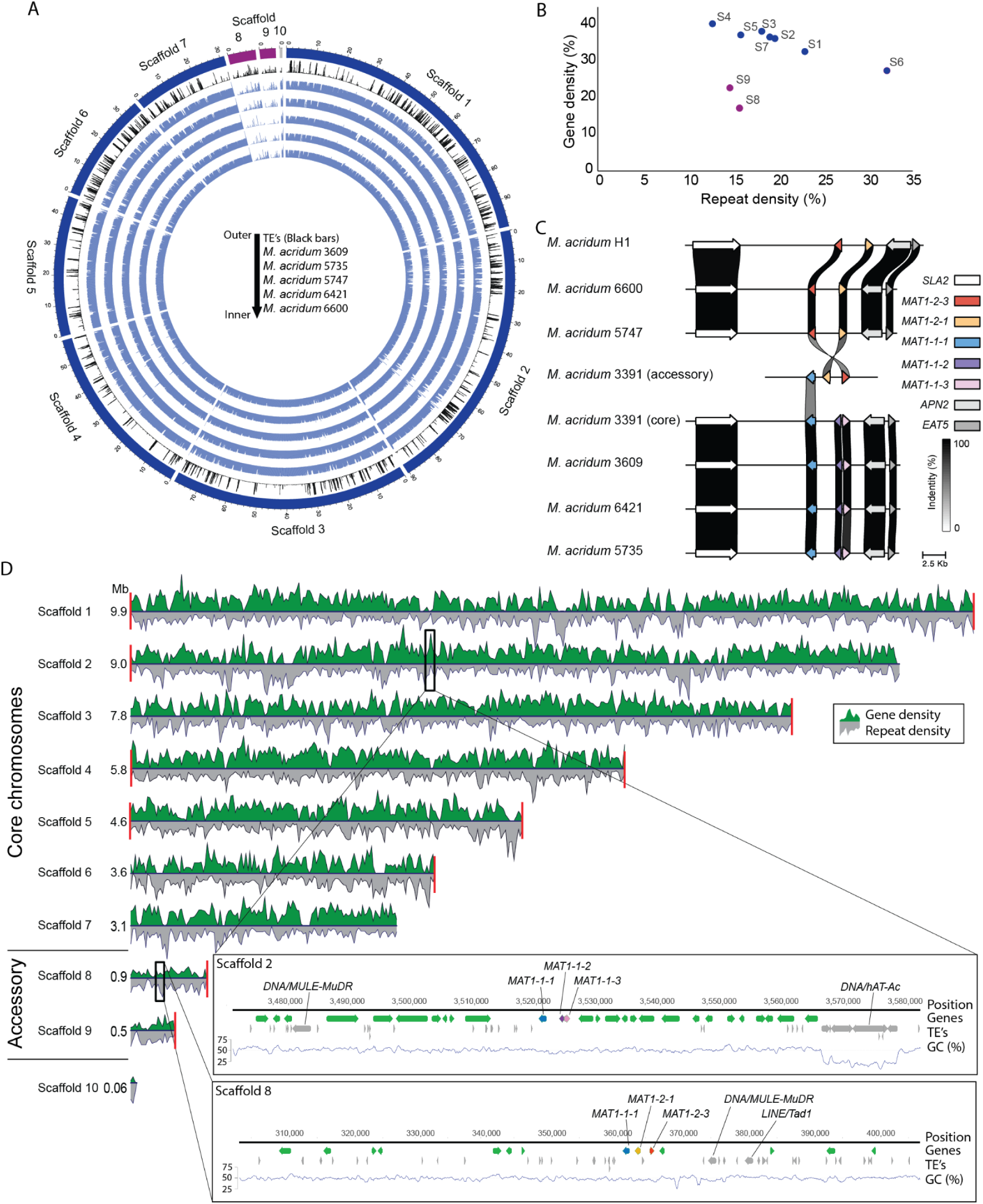
Accessory chromosome with MAT locus in *M. acridum.* (A) Circos plot showing the 10 scaffolds of ARSEF 3391 in the outermost dark blue (core chromosomes) and purple (accessory chromosomes) rings. Identified transposable elements (TE’s) are shown as black bars followed by short-read coverage of each of the five other *M. acridum* isolates mapped on to ARSEF 3391 showing that scaffold 8 and 9 are unique to ARSEF 3391. (B) Gene density vs repeat density plotted for each scaffold of ARSEF 3391. Blue dots are core chromosomes (scaffolds S1-7) and purple dots are scaffold 8 and 9 making up the accessory chromosome. (C) Synteny of the canonical mating type (*MAT*) locus among isolates. The *MAT* locus flanking genes, *SLA2* and *APN2*, were highly conserved in both *MAT1-1* and *MAT1-2 M. acridum* individuals. Four of the six isolates possessed the *MAT1-1* idiomorph (*MAT1-1-1, MAT1-1-2*, and *MAT1-1-3*), while two of the six as well as the previously sequenced reference strain possessed the *MAT1-2* idiomorph (*MAT1-2-1* and *MAT1-2-3*). ARSEF 3391 additionally harboured *MAT*-associated sequences on its accessory chromosome, possessing a slightly truncated, but likely still functional *MAT1-1-1* and as intact *MAT1-2-1* and *MAT1-2-3* genes. (D) Genome structure of ARSEF 3391 showing the gene density and repeat density along the 10 scaffolds. Red bars at the end of a scaffold indicate telomeric repeats. The two inserted boxes show placement and genomic regions of the *MAT* loci identified on the core chromosome (scaffold 2) and accessory chromosome (scaffold 8), respectively.

Careful manual annotation of mating-type loci in *M. acridum* revealed that all isolates contained either a highly conserved canonical *MAT1-1* or *MAT1-2* idiomorph (**Fig. 5c**), as expected for this heterothallic species [42]. These loci were flanked by the *SLA2* and *APN2* genes as is typical for Sordariomycete fungi. In each case, these genes encoded predicted proteins harbouring their respective functional domains: MAT1-1-1 (MATalpha_HMGbox, cl07856), MAT1-1-3 (HMG-box_ROX1-like, cd01389), and MAT1-2-1 (HMG-box_ROX1-like, cd01389). Additionally, the serine residues within the HMG box domains were split by an intron, the defining feature of the MAT-associated HMG box domain. Furthermore, in addition to the *MAT1-1* idiomorph on core scaffold 2, isolate ARSEF 3391 contained a second mating-type locus consisting of less conserved *MAT1-1-1, MAT1-2-1,* and *MAT1-2-3* genes, as well as a heavily pseudogenised version of the *SLA2* gene loci on the accessory chromosome (**Fig. 5c-d**).

When used as queries against the NCBI NR databases, the accessory *MAT1-1-1* and *MAT1-2-1* genes from *M. acridum* ARSEF 3391 were more similar to other *Metarhizium* species than *M. acridum* (**Table S8-S11**). The subsequent phylogenetic analyses supported this result, illustrating that the *MAT1-1-1*, *MAT1-2-1* and *MAT1-2-3* genes from the accessory chromosome of *M. acridum* ARSEF 3391 did not cluster with the *M. acridum* homologs (**Fig. S16-S19**). Despite the phylogenetic distance suggesting that this putative *MAT* loci on the accessory chromosome may have been horizontally acquired from a different *Metarhizium* species, the locus may still be functional and active because at least *MAT1-2-3* is expressed to similar levels as genes in the core *MAT* locus (**Fig. S20**).

To assess if the accessory chromosome also contains virulence related genes as seen on accessory chromosomes of plant pathogenic fungi, we compared the distribution of functional gene groups between accessory and core chromosomes on the ARSEF 3391 genome (**Fig. 6a-h**). However, this *M. acridum* accessory chromosome does not contain any effectors or secondary metabolite synthesis genes (**Fig. 6b,d**). Further analyzing the 215 genes on scaffold 8 and 9 that make up this accessory chromosome showed the majority of annotated genes to be singletons or accessory in relation to *M. acridum* pangenome (**Fig. 6i**). 20.3 % of the genes on the accessory chromosome were part of the core *M. acridum* pangenome gene content, but this is because they have paralogs or xenologs in other isolates and therefore are designated as core.

**Fig. 6:**
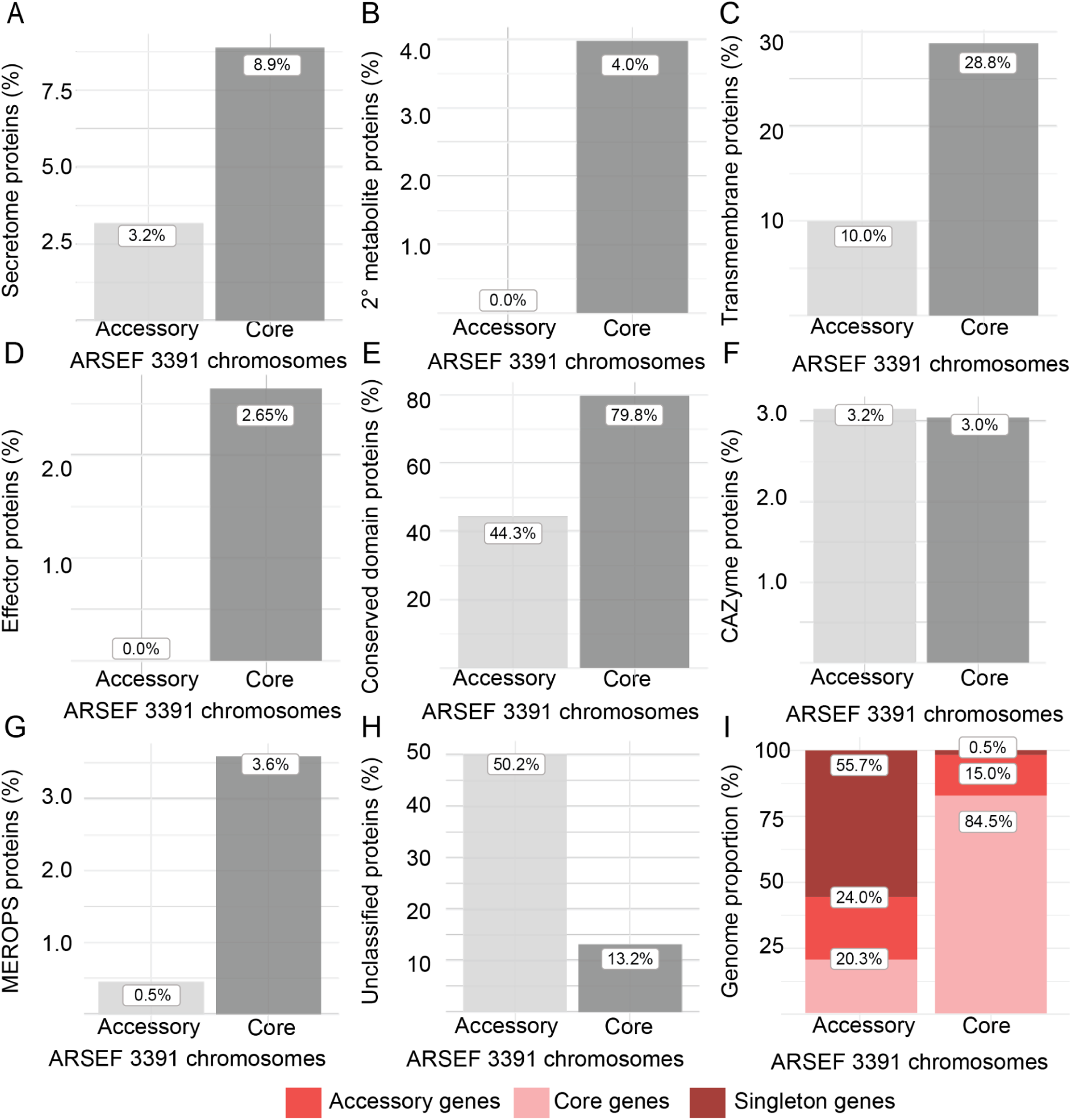
Accessory chromosome gene content. Percentage of genes on accessory vs. core chromosomes that are functionally assigned as secretome proteins (A), Secondary metabolite genes (B), Transmembrane proteins (C), Effector proteins (D), conserved domain proteins (E), CAZymes (F), MEROPS proteases (G), or unclassified proteins (H). (I) The percentage of genes assigned as either core genes, accessory genes, or singletons on either accessory chromosomes or core chromosomes.

## DISCUSSION

Pathogenic fungi undergo rapid evolution in response to changing environments and host immune measures. The division of the genome into gene-dense core compartments—harboring essential genes protected from high mutation rates, transposable elements, and novel gene insertions—and gene-sparse dynamic regions allows pathogenic fungi to balance genome stability with rapid adaptability to hosts and changing environments [4, 80]. Here we show that the intra-specific pangenome of the entomopathogenic fungus, *M. acridum,* is compartmentalized into gene-sparse and gene-rich regions and has undergone extensive genome restructuring. We find that the pangenome of *M. acridum* contains a larger gene repertoire than reported from two previously sequenced genomes [42, 78], with almost 25% of the gene content composed of singleton or accessory genes, which are thus dispensable.

Genes in the accessory compartment of the genome have a higher percentage of effectors and together with insect cuticle degrading serine proteases (Peptidase S8) and chitinases (Glycoside Hydrolase family 18), effectors are enriched in TE-rich gene-sparse regions of the *M. acridum* genome. The content and distribution of accessory genes within pangenomes are structured and maintained by selection [81, 82], and within the genus *Metarhizium,* serine proteases and chitinases are positively selected and rapidly evolving [26]. The fungus *M. acridum* has been shown to contain the highest number of rapidly evolving positively selected genes within the genus [26]. Studies in other taxa have shown similar patterns, with directional selection playing a major role in the acclimatization of a pathogen to the environment of a novel host [83–85], which is consistent with accessory loci associated with host-pathogen traits directly affecting population differentiation and host niche adaptation in *M. acridum*.

The genus, *Metarhizium*, is a diverse clade of entomopathogenic fungi with host ranges occurring across the spectrum from generalist to specialist [86]. Inter-species comparative genomics and genetics have shown major differences in the gene dynamics of *Metarhizium* species [26, 28, 29, 78]. Here, we identify a total of 17,106 orthogroups in the genus, with 5,136 (36.9%) forming the core orthogroups of *Metarhizium*. We observe evidence for a large duplication event at the node separating generalists from specialists (103 orthogroups; **Fig. S4**). This is in accordance with previous studies that have shown high numbers of paralogs in generalists, likely due to the lack of active RIP mechanisms to prevent gene duplication in such species [26]. However, we also see evidence for gene duplication in specialists, with 41 duplicated orthologs in the node that splits isolates of *M. acridum* from each other. Gene duplication events are a powerful source of functional innovation [87]. We already see evidence of such duplications in the ARSEF 324, a unique isolate of *M. acridum* that is shown to be functionally diploid [42].

Our investigation of *M. acridum* revealed 739 orthogroups that were exclusive to the species. Of these, 255 protein clusters formed a core subset specific to *M. acridum* that was surpassed only by the number of species-specific orthogroups in *M. rileyi* (343; **Fig. 2b**). Further analysis of these 255 orthogroups showed that they were enriched in gene functions critical to host-pathogen interactions, such as the synthesis of secondary metabolites and serine proteases. Fungi use secondary metabolites as a chemical arsenal to improve their fitness in challenging environments, such as the hazardous environment of the host body during cuticle penetration and infection of the hemocoel [88]. *Metarhizium* specialist species, such as *M. acridum*, are known to display increased host immune evasion tactics compared to their generalist counterparts [89]. Therefore, the essential core subset of proteins unique to *M. acridum* may have arisen as necessary adaptations for specializing to their orthopteran hosts and for launching subversive infections while avoiding immune defense responses. Large-scale bacterial pangenome studies have shown that the size of a pangenome is strongly correlated to niche heterogeneity, diverse species interactions, and the effective population size of a species [90, 91]. Given its widespread distribution and potential to infect over 26,000 extant orthopteran host species [45], the presence of accessory genome elements in *M. acridum* may serve as a crucial component of the pathogen’s adaptive strategy to colonize and thrive in a diverse range of ecological niches and hosts.

In some fungi, genome compartmentalization extends to include entire accessory chromosomes [20], and within the genus *Metarhizium, M. robertsii* and *M. guizhouense* have been shown to contain horizontally transferred accessory chromosomes [75, 77]. Here we identified an ∼1.4 Mb accessory chromosome in *M. acridum*, which is only present in isolate ARSEF 3391 from Tanzania. In fungi, accessory chromosomes typically exhibit similar structural characteristics, ranging from 0.2-3.5 Mb in length, being enriched in transposons and repetitive sequences, having lower gene density, or exhibiting alternative codon usage in comparison to the core genome [22, 92–94]. This is also the case for accessory chromosomes in *M. robertsii,* which are enriched in transposable elements and contain effector proteins similar to our identified gene-sparse regions in *M. acridum* [75]. However, we did not identify any effectors on either of the two scaffolds 8 and 9 in ARSEF 3391 that together make up the accessory chromosome. The accessory chromosome in *M. acridum* does not contain a higher percentage of transposons or repetitive regions compared to core chromosomes, but does contain a markedly different TE landscape (**Fig. S14**). The accessory chromosome contains a higher proportion of DNA/MULE elements and markedly higher proportions of LINE elements. These TE elements are nearly absent in core chromosomes that instead show the typical ascomycete pattern of ca. 50% LTRs. Such markedly different TE landscapes are consistent with different inheritance patterns and a potential past horizontal chromosome transfer [20], indicating different evolutionary histories of the core chromosomes and this accessory chromosome.

The activity of Class II (DNA) transposons may influence the general genome organization in *M. acridum*. Similar to what has been found in *M. anisopliae* [77], the activity of three different DNA transposons—hAT-Ac, Helitron, and MULE-MuDR—also appears to play a role in genome organization in *M. acridum*. These elements could contribute to genome reshuffling by inducing double-or single-stranded breaks, or alternatively, by providing substrates for non-homologous recombination; however, the exact mechanism remains unclear. Interestingly, the expansion of DNA transposons is most pronounced in *M. acridum* ARSEF 3391, which possesses an accessory chromosome. Indeed, all three DNA transposons also have highly similar copies on this accessory chromosome (scaffolds 8 and 9). It is unclear, however, whether this accessory chromosome serves as the source of these DNA transposons or as a target, or whether these TEs were acquired horizontally through a different mechanism—for example, as described for *M. anisopliae* via the giant Starship transposons [17, 21, 77, 95, 96].

In fungi, accessory chromosomes are thought to confer a fitness effect in specific environments [97, 98], exemplified by the fungal wheat pathogen, *Zymoseptoria tritici*, where the presence of certain accessory chromosomes affects pathogenicity against a specific wheat cultivar [99]. In our functional enrichment analysis of accessory chromosomes in ARSEF 3391, we found enriched functions primarily correlated with metabolism, but we also discovered several enriched functions related to sexual reproduction. Unlike many *Metarhizium* species, which are known only from their asexual morphs and show limited evidence of recombination in nature [32]*, M. acridum* is also known only from its asexual form but shows genomic signs of sexual reproduction, primarily through the presence of RIP mutations [42]. Detailed annotation of mating-type (*MAT*) loci showed the mutually exclusive presence of *MAT1-1* and *MAT1-2* mating-type idiomorphs among our six isolates, consistent with outcrossing heterothallism and which has also been determined within the *M. anisopliae-*group [100]. However, on the accessory chromosome of ARSEF 3391, we identified a non-canonical *MAT* locus, lacking the surrounding *SLA2, APN2, and EAT5* genes often used to identify *MAT* loci but having the full complementary *MAT* locus, including the *MAT1-2-1,* and *MAT1-2-3* genes, in addition to *MAT1-1-1.* Phylogenetic analysis of the *M. acridum MAT* loci show that genes on the accessory chromosome *MAT* locus are related to other *Metarhizium* species, rather than *M. acridum*, clustering with early diverging lineages such as *Pochonia chlamydospora* and *M. rileyi* within the *Metarhizium* lineage [41]. The isolate thus harbors the *MAT1-1* locus on a core chromosome, along with genes from the *MAT1-2* locus located on an accessory chromosome. If this non-canonical accessory mating-type locus is functional, the isolate ARSEF 3391 may therefore be effectively homothallic. This is supported by the observation that the *MAT1-2* locus on the accessory chromosome is actively expressed, with the *MAT1-2-3* gene showing expression levels comparable to the *MAT1-1* locus on the core chromosome.

We searched for *captain* elements (DUF3435 tyrosine recombinase) in the vicinity of the accessory *MAT* locus to determine if a *starship* element might have carried the *MAT* locus onto the accessory chromosome similar to a recently identified TE expansion in a *M. anisopliae* isolate [77]. However, this does not appear to be the case since the nearest *captain* domain is located ca. 50 kb upstream, on the plus strand and thus orienting the putative *starship* element away from the accessory *MAT* locus. Accessory chromosomes contribute significantly to genomic diversity in many fungi, but may randomly be lost during cell division and growth. While some accessory chromosomes may be maintained because they provide a selective advantage during fungal colonization of hosts, some accessory chromosomes appear to be maintained without providing a benefit to the fungus. It has thus been speculated that accessory chromosomes carry a signal that contributes to their preferential transfer and maintenance during cell divisions and meiosis [20]. In some fungi such as *Podospora* and *Fusarium,* accessory chromosomes carry genes homologous to meiotic drive toxin/antidote proteins (*Spoks*), which could explain the continued maintenance of fungal cells carrying accessory chromosomes compared to cells without accessory chromosomes [101]. The fact that we find a *MAT* locus an accessory chromosome in *M. acridum* potentially suggests a different mechanism for maintenance and effect of accessory chromosomes to the fungal host. Having the accessory chromosomes would then remove the necessity of finding a mating partner of the opposite mating type. Furthermore, since the non-canonical *MAT* locus carry genes of both the *MAT1-1* and *MAT1-2* idiomorph, this could theoretically contribute to the maintenance and preferential transfer of this accessory chromosome, although the exact mechanism is unknown.

For many species of *Metarhizium* the sexual stage has never been observed and these fungi are considered facultatively sexual, or exclusively asexual. This is also evident for some of the most widespread and abundant species such as *M. robertsii* and *M. anisopliae*, population structure is primarily clonal and sexual reproduction appears to be very rare [32, 100]. The obligate locust pathogen studied here, *M. acridum,* also primarily reproduces clonally but show signs of more frequent recombination [42]. Without frequent meiotic recombination, genome structural plasticity may provide an alternative to facilitate asexual genome evolution. Genomes of fungal pathogens are known to be highly dynamic [93], and we observe large-scale chromosomal translocations and inversions in *M. acridum.* With organisms that are primarily asexually propagating, intra- and inter-chromosomal rearrangements are thought to be a significant driver of genetic variation [92, 94]. However, the extensive non-synteny among most of the isolates studied here suggests that their chromosomes are too divergent for homologous pairing and recombination, and possibly represents beginning speciation. Whether the capability for sexual reproduction between locally occurring isolates with more homologous chromosomal structure is facilitated by the presence of accessory chromosomes remains to be investigated.

## ACKOWLEDGMENTS

This research was supported by grants from the Independent Research Fund Denmark Starting Grant (No. 8049-00086B) awarded to H. H. De Fine Licht and from the European Union’s Horizon 2020 research and innovation programme under the Marie SklodowskaCurie grant agreement No. 801199.

## REFERENCES

1. Möller M, Stukenbrock EH. Evolution and genome architecture in fungal plant pathogens. Nat Rev Microbiol 2017;15:756–771. 10.1038/nrmicro.2017.76

2. Wacker T et al. Genome variation in the Batrachochytrium pathogens of amphibians. PLOS Pathog 2024;20:e1012218. 10.1371/journal.ppat.1012218

3. Wacker T et al. Two-speed genome evolution drives pathogenicity in fungal pathogens of animals. Proc Natl Acad Sci 2023;120:e2212633120. 10.1073/pnas.2212633120

4. Torres DE et al. Genome evolution in fungal plant pathogens: looking beyond the two-speed genome model. Fungal Biol Rev 2020;34:136–143. 10.1016/j.fbr.2020.07.001

5. Sánchez-Vallet A et al. The Genome Biology of Effector Gene Evolution in Filamentous Plant Pathogens. Annu Rev Phytopathol 2018;56:21–40. 10.1146/annurev-phyto-080516-035303

6. Croll D. Dimensions of genome dynamics in fungal pathogens: from fundamentals to applications. BMC Biol 2024;22:19. 10.1186/s12915-023-01786-w

7. Tettelin H et al. Genome analysis of multiple pathogenic isolates of *Streptococcus agalactiae* : Implications for the microbial “pan-genome”. Proc Natl Acad Sci 2005;102:13950–13955. 10.1073/pnas.0506758102

8. Wang K et al. The Chicken Pan-Genome Reveals Gene Content Variation and a Promoter Region Deletion in *IGF2BP1* Affecting Body Size. Mol Biol Evol 2021;38:5066–5081. 10.1093/molbev/msab231

9. Zhou Y et al. Assembly of a pangenome for global cattle reveals missing sequences and novel structural variations, providing new insights into their diversity and evolutionary history. Genome Res 2022;32:1585–1601. 10.1101/gr.276550.122

10. Badet T et al. A 19-isolate reference-quality global pangenome for the fungal wheat pathogen Zymoseptoria tritici. BMC Biol 2020;18:12. 10.1186/s12915-020-0744-3

11. McCarthy CGP, Fitzpatrick DA. Pan-genome analyses of model fungal species. Microb Genomics 2019;5. 10.1099/mgen.0.000243

12. Perrier M, Barber AE. Unraveling the genomic diversity and virulence of human fungal pathogens through pangenomics. PLOS Pathog 2024;20:e1012313. 10.1371/journal.ppat.1012313

13. Torres DE, Thomma BPHJ, Seidl MF. Transposable Elements Contribute to Genome Dynamics and Gene Expression Variation in the Fungal Plant Pathogen *Verticillium dahliae*. Genome Biol Evol 2021;13:evab135. 10.1093/gbe/evab135

14. Plissonneau C, Hartmann FE, Croll D. Pangenome analyses of the wheat pathogen Zymoseptoria tritici reveal the structural basis of a highly plastic eukaryotic genome. BMC Biol 2018;16:5. 10.1186/s12915-017-0457-4

15. Hayward A, Gilbert C. Transposable elements. Curr Biol 2022;32:R904–R909. 10.1016/j.cub.2022.07.044

16. Díaz CL et al. Transposons and accessory genes drive adaptation in a clonally evolving fungal pathogen. 2025. Microbiology, 2025.

17. Urquhart A, Vogan AA, Gluck-Thaler E. Starships: a new frontier for fungal biology. Trends Genet 2024;40:1060–1073. 10.1016/j.tig.2024.08.006

18. Urquhart AS, Gluck-Thaler E, Vogan AA. Gene acquisition by giant transposons primes eukaryotes for rapid evolution via horizontal gene transfer. Sci Adv 2024.

19. Urquhart AS et al. *Starships* are active eukaryotic transposable elements mobilized by a new family of tyrosine recombinases. Proc Natl Acad Sci 2023;120:e2214521120. 10.1073/pnas.2214521120

20. Habig M et al. Horizontal transfer of accessory chromosomes in fungi – a regulated process for exchange of genetic material? Heredity 2025. 10.1038/s41437-025-00746-0

21. Urquhart AS et al. A natural mechanism of eukaryotic horizontal gene transfer. 2025. Evolutionary Biology, 2025.

22. Ma L-J et al. Comparative genomics reveals mobile pathogenicity chromosomes in Fusarium. Nature 2010;464:367–373. 10.1038/nature08850

23. Metzenberg RL, Glass NL. Mating type and mating strategies in *Neurospora*. BioEssays 1990;12:53–59. 10.1002/bies.950120202

24. Wilken PM et al. Which MAT gene? Pezizomycotina (Ascomycota) mating-type gene nomenclature reconsidered. Fungal Biol Rev 2017;31:199–211. 10.1016/j.fbr.2017.05.003

25. Whittle CA, Nygren K, Johannesson H. Consequences of reproductive mode on genome evolution in fungi. Fungal Genet Biol 2011;48:661–667. 10.1016/j.fgb.2011.02.005

26. Hu X et al. Trajectory and genomic determinants of fungal-pathogen speciation and host adaptation. Proc Natl Acad Sci 2014;111:16796–16801. 10.1073/pnas.1412662111

27. Joop G, Vilcinskas A. Coevolution of parasitic fungi and insect hosts. Zoology 2016;119:350–358. 10.1016/j.zool.2016.06.005

28. Zhang Q et al. Horizontal gene transfer allowed the emergence of broad host range entomopathogens. Proc Natl Acad Sci 2019;116:7982–7989. 10.1073/pnas.1816430116

29. Gao B-J et al. Subtilisin-like Pr1 proteases marking the evolution of pathogenicity in a wide-spectrum insect-pathogenic fungus. Virulence 2020;11:365–380. 10.1080/21505594.2020.1749487

30. Stone LBL, Bidochka MJ. The multifunctional lifestyles of Metarhizium: evolution and applications. Appl Microbiol Biotechnol 2020;104:9935–9945. 10.1007/s00253-020-10968-3

31. Driver F, Milner RJ, Trueman JWH. A taxonomic revision of Metarhizium based on a phylogenetic analysis of rDNA sequence data. Mycol Res 2000;104:134–150. 10.1017/S0953756299001756

32. Rehner SA. Genetic structure of Metarhizium species in western USA: Finite populations composed of divergent clonal lineages with limited evidence for recent recombination. J Invertebr Pathol 2020;177:107491. 10.1016/j.jip.2020.107491

33. Moonjely S, Bidochka MJ. Generalist and specialist Metarhizium insect pathogens retain ancestral ability to colonize plant roots. Fungal Ecol 2019;41:209–217. 10.1016/j.funeco.2019.06.004

34. Milner RJ et al. A comparative study of two Mexican isolates with an Australian isolate of Metarhizium anisopliae var. acridum – strain characterisation, temperature profile and virulence for wingless grasshopper, Phaulacridium vittatum.

35. Brunner-Mendoza C et al. A review on the genus *Metarhizium* as an entomopathogenic microbial biocontrol agent with emphasis on its use and utility in Mexico. Biocontrol Sci Technol 2019;29:83–102. 10.1080/09583157.2018.1531111

36. Li Z et al. Biological control of insects in Brazil and China: history, current programs and reasons for their successes using entomopathogenic fungi. Biocontrol Sci Technol 2010;20:117–136. 10.1080/09583150903431665

37. Aw KMS, Hue SM. Mode of Infection of Metarhizium spp. Fungus and Their Potential as Biological Control Agents. J Fungi 2017;3:30. 10.3390/jof3020030

38. Wang C, St. Leger RJ. Developmental and Transcriptional Responses to Host and Nonhost Cuticles by the Specific Locust Pathogen *Metarhizium anisopliae* var. *acridum*. Eukaryot Cell 2005;4:937–947. 10.1128/EC.4.5.937-947.2005

39. Hunter DM. Credibility of an IPM Approach for Locust and Grasshopper Control: The Australian Example*. J Orthoptera Res 2010;19:133–137. 10.1665/034.019.0108

40. Fernandes ÃKK et al. Characterization of Metarhizium species and varieties based on molecular analysis, heat tolerance and cold activity: Metarhizium: variation in DNA and temperature tolerances. J Appl Microbiol 2010;108:115–128. 10.1111/j.1365-2672.2009.04422.x

41. Mongkolsamrit S et al. Revisiting Metarhizium and the description of new species from Thailand. Stud Mycol 2020;95:171–251. 10.1016/j.simyco.2020.04.001

42. Nielsen KN et al. Diploidy within a Haploid Genus of Entomopathogenic Fungi. Genome Biol Evol 2021;13:evab158. 10.1093/gbe/evab158

43. Parker D, Meyling NV, De Fine Licht HH. Phenotypic variation and genomic variation in insect virulence traits reveal patterns of intraspecific diversity in a locust-specific fungal pathogen. J Evol Biol 2023;jeb.14214. 10.1111/jeb.14214

44. Cigliano MM et al. Orthoptera Species File. Version 5.0/5.0. http://Orthoptera.SpeciesFile.org. http://Orthoptera.SpeciesFile.org. (November 2022, date last accessed).

45. Song H. Biodiversity of Orthoptera. Insect Biodiversity: Science and Society, 1st edn. Wiley, 2018, 245–279.

46. Saud Z et al. Telomere length de novo assembly of all 7 chromosomes and mitogenome sequencing of the model entomopathogenic fungus, Metarhizium brunneum, by means of a novel assembly pipeline. BMC Genomics 2021;22:87. 10.1186/s12864-021-07390-y

47. Li H. Minimap2: pairwise alignment for nucleotide sequences. Bioinformatics 2018;34:3094–3100. 10.1093/bioinformatics/bty191

48. Kolmogorov M et al. Assembly of long, error-prone reads using repeat graphs. Nat Biotechnol 2019;37:540–546. 10.1038/s41587-019-0072-8

49. Chen Y et al. Efficient assembly of nanopore reads via highly accurate and intact error correction. Nat Commun 2021;12:60. 10.1038/s41467-020-20236-7

50. Gurevich A et al. QUAST: quality assessment tool for genome assemblies. Bioinformatics 2013;29:1072–1075. 10.1093/bioinformatics/btt086

51. Jackman SD et al. Tigmint: correcting assembly errors using linked reads from large molecules. BMC Bioinformatics 2018;19:393. 10.1186/s12859-018-2425-6

52. Baril T, Galbraith J, Hayward A. Earl Grey: A Fully Automated User-Friendly Transposable Element Annotation and Analysis Pipeline. Mol Biol Evol 2024;41:msae068. 10.1093/molbev/msae068

53. Gluck-Thaler E, Vogan AA. Systematic identification of cargo-mobilizing genetic elements reveals new dimensions of eukaryotic diversity. Nucleic Acids Res 2024;52:5496–5513. 10.1093/nar/gkae327

54. Jonathan M. Palmer, Jason Stajich. Funannotate v1.8.1: Eukaryotic genome annotation. 2020. 2020.

55. Slater GSC, Birney E. Automated generation of heuristics for biological sequence comparison. BMC Bioinformatics 2005;6:31. 10.1186/1471-2105-6-31

56. Lowe TM, Eddy SR. tRNAscan-SE: a program for improved detection of transfer RNA genes in genomic sequence. Nucleic Acids Res 1997;25.

57. Haas BJ et al. Automated eukaryotic gene structure annotation using EVidenceModeler and the Program to Assemble Spliced Alignments. Genome Biol 2008;9:R7. 10.1186/gb-2008-9-1-r7

58. Apweiler R. UniProt: the Universal Protein knowledgebase. Nucleic Acids Res 2004;32:115D – 119. 10.1093/nar/gkh131

59. Jones P et al. InterProScan 5: genome-scale protein function classification. Bioinformatics 2014;30:1236–1240. 10.1093/bioinformatics/btu031

60. Almagro Armenteros JJ et al. SignalP 5.0 improves signal peptide predictions using deep neural networks. Nat Biotechnol 2019;37:420–423. 10.1038/s41587-019-0036-z

61. Käll L, Krogh A, Sonnhammer ELL. A Combined Transmembrane Topology and Signal Peptide Prediction Method. J Mol Biol 2004;338:1027–1036. 10.1016/j.jmb.2004.03.016

62. Finn RD, Clements J, Eddy SR. HMMER web server: interactive sequence similarity searching.

63. Zhang H et al. dbCAN2: a meta server for automated carbohydrate-active enzyme annotation. Nucleic Acids Res 2018;46:W95–W101. 10.1093/nar/gky418

64. Rawlings ND, Barrett AJ, Finn R. Twenty years of the *MEROPS* database of proteolytic enzymes, their substrates and inhibitors. Nucleic Acids Res 2016;44:D343–D350. 10.1093/nar/gkv1118

65. Blin K et al. antiSMASH 5.0: updates to the secondary metabolite genome mining pipeline. Nucleic Acids Res 2019;47:W81–W87. 10.1093/nar/gkz310

66. Simão FA et al. BUSCO: assessing genome assembly and annotation completeness with single-copy orthologs. Bioinformatics 2015;31:3210–3212. 10.1093/bioinformatics/btv351

67. Emms DM. OrthoFinder: solving fundamental biases in whole genome comparisons dramatically improves orthogroup inference accuracy.

68. Emms DM. OrthoFinder: phylogenetic orthology inference for comparative genomics.

69. Emms DM, Kelly S. STAG: Species Tree Inference from All Genes. 2018. 10.1101/267914

70. Emms DM, Kelly S. STRIDE: Species Tree Root Inference from Gene Duplication Events.

71. Falcon S, Gentleman R. Using GOstats to test gene lists for GO term association. Bioinformatics 2007;23:257–258. 10.1093/bioinformatics/btl567

72. Lovell JT et al. GENESPACE tracks regions of interest and gene copy number variation across multiple genomes. eLife 2022;11:e78526. 10.7554/eLife.78526

73. Quinlan AR, Hall IM. BEDTools: a flexible suite of utilities for comparing genomic features. Bioinformatics 2010;26:841–842. 10.1093/bioinformatics/btq033

74. Li H, Durbin R. Fast and accurate long-read alignment with Burrows–Wheeler transform. Bioinformatics 2010;26:589–595. 10.1093/bioinformatics/btp698

75. Habig M et al. Frequent horizontal chromosome transfer between asexual fungal insect pathogens. Proc Natl Acad Sci 2024;121. 10.1073/pnas.2316284121

76. Van Den Belt M et al. CAGECAT: The CompArative GEne Cluster Analysis Toolbox for rapid search and visualisation of homologous gene clusters. BMC Bioinformatics 2023;24:181. 10.1186/s12859-023-05311-2

77. Griem-Krey H et al. Transposable elements hitchhike on Starships across fungal genomes.

78. Gao Q et al. Genome Sequencing and Comparative Transcriptomics of the Model Entomopathogenic Fungi Metarhizium anisopliae and M. acridum. PLoS Genet 2011;7:e1001264. 10.1371/journal.pgen.1001264

79. Muszewska A et al. Fungal lifestyle reflected in serine protease repertoire. Sci Rep 2017;7:9147. 10.1038/s41598-017-09644-w

80. Croll D, McDonald BA. The Accessory Genome as a Cradle for Adaptive Evolution in Pathogens. PLoS Pathog 2012;8:e1002608. 10.1371/journal.ppat.1002608

81. Goyal A. Metabolic adaptations underlying genome flexibility in prokaryotes. PLOS Genet 2018;14:e1007763. 10.1371/journal.pgen.1007763

82. Whelan FJ, Hall RJ, McInerney JO. Evidence for Selection in the Abundant Accessory Gene Content of a Prokaryote Pangenome. Mol Biol Evol 2021;38:3697–3708. 10.1093/molbev/msab139

83. Zhong Z et al. Directional Selection from Host Plants Is a Major Force Driving Host Specificity in Magnaporthe Species. Sci Rep 2016;6:25591. 10.1038/srep25591

84. Raffaele S et al. Genome Evolution Following Host Jumps in the Irish Potato Famine Pathogen Lineage. Science 2010;330:1540–1543. 10.1126/science.1193070

85. Stukenbrock EH, Bataillon T. A Population Genomics Perspective on the Emergence and Adaptation of New Plant Pathogens in Agro-Ecosystems. PLoS Pathog 2012;8:e1002893. 10.1371/journal.ppat.1002893

86. St. Leger RJ, Wang JB. *Metarhizium* : jack of all trades, master of many. Open Biol 2020;10:200307. 10.1098/rsob.200307

87. Wapinski I et al. Natural history and evolutionary principles of gene duplication in fungi. Nature 2007;449:54–61. 10.1038/nature06107

88. Rohlfs M, Churchill ACL. Fungal secondary metabolites as modulators of interactions with insects and other arthropods. Fungal Genet Biol 2011;48:23–34. 10.1016/j.fgb.2010.08.008

89. Li J, Xia Y. Host–Pathogen Interactions between Metarhizium spp. and Locusts. J Fungi 2022;8:602. 10.3390/jof8060602

90. Rouli L et al. The bacterial pangenome as a new tool for analysing pathogenic bacteria. New Microbes New Infect 2015;7:72–85. 10.1016/j.nmni.2015.06.005

91. Segerman B. The genetic integrity of bacterial species: the core genome and the accessory genome, two different stories. Front Cell Infect Microbiol 2012;2. 10.3389/fcimb.2012.00116

92. Habig M, Stukenbrock EH. 2 Origin, Function, and Transmission of Accessory Chromosomes. Genetics and Biotechnology. Cham: Springer International Publishing, 2020, 25–47.

93. Goodwin SB et al. Finished Genome of the Fungal Wheat Pathogen Mycosphaerellagraminicola Reveals Dispensome Structure, Chromosome Plasticity, and Stealth Pathogenesis.

94. Coleman JJ et al. The Genome of Nectria haematococca: Contribution of Supernumerary Chromosomes to Gene Expansion. PLoS Genet 2009;5:e1000618. 10.1371/journal.pgen.1000618

95. Gluck-Thaler E et al. Giant transposons promote strain heterogeneity in a major fungal pathogen. 2025. Microbiology, 2025.

96. Gluck-Thaler E et al. Giant *Starship* Elements Mobilize Accessory Genes in Fungal Genomes. Mol Biol Evol 2022;39:msac109. 10.1093/molbev/msac109

97. Bertazzoni S et al. Accessories Make the Outfit: Accessory Chromosomes and Other Dispensable DNA Regions in Plant-Pathogenic Fungi. Mol Plant-Microbe Interactions® 2018;31:779–788. 10.1094/MPMI-06-17-0135-FI

98. Witte TE et al. Accessory Chromosome-Acquired Secondary Metabolism in Plant Pathogenic Fungi: The Evolution of Biotrophs Into Host-Specific Pathogens. Front Microbiol 2021;12. 10.3389/fmicb.2021.664276

99. Habig M, Quade J, Stukenbrock EH. Forward Genetics Approach Reveals Host Genotype-Dependent Importance of Accessory Chromosomes in the Fungal Wheat Pathogen *Zymoseptoria tritici*. mBio 2017;8:e01919–17. 10.1128/mBio.01919-17

100. Rehner SA, Kepler RM. Species limits, phylogeography and reproductive mode in the Metarhizium anisopliae complex. J Invertebr Pathol 2017;148:60–66. 10.1016/j.jip.2017.05.008

101. Sandell L et al. The role of toxin/antidote genes in the maintenance and evolution of accessory chromosomes in*Fusarium*. 2025. Cold Spring Harbor Laboratory, 2025.

